# Chemically induced dislocation and mutation of essential cytoskeletal proteins: A case study focusing on Coronin-A

**DOI:** 10.1101/2024.08.05.606068

**Authors:** Maike B. Stange, Robert Großmann, Sven Flemming, Markus Grebe, Carsten Beta

**Affiliations:** Institute of Physics and Astronomy, University of Potsdam, D-14476 Potsdam, Germany; Institute of Biochemistry and Biology, University of Potsdam, D-14476 Potsdam, Germany

## Abstract

Actin is one of the most highly conserved proteins—the actin cytoskeleton or actin-like structures are found in all kingdoms of life. In this work, Coronin-A, a central player involved in actin dynamics, is considered as an example to compare the effect of four mutation strategies on its function in *Dictyostelium discoideum*. Two distinct genetic knockout methods (selection-linked integration and classical gene disruption), RNAi knockdown, and sudden loss-of-function generated by chemically induced dislocation (CID) are compared and the cell lines are characterized according to actin dependent processes: appearance, development, and motility. These phenotypic features are affected to different degrees in the mutant cell lines, providing a basis to discuss the strengths and weaknesses of each approach. This comparative study highlights the advantages of the newly introduced selection-linked integration method to knock out genes over commonly used gene disruption, and for expressing a CID-based knock-sideways system under the native promotor of the protein of interest.

## INTRODUCTION

The actin cytoskeleton is a major structural element in eukaryotic cells that determines their shape and mechanics. It forms a branched cortical fiber network that is essential for maintaining cell shape and for stabilizing the nucleus. Its dynamical properties allow rapid reorganization of the filament network and drive important cellular functions, such as migration, nutrient uptake, rearrangement of intracellular components, and growth. The dynamical character of the actin cytoskeleton is maintained by a so-called treadmilling mechanism, in which actin fibers grow by the addition of ATP-bound monomers at one end, while less active ADP-bound monomers are removed from the other end [1–3]. Together with a large number of auxiliary proteins that, for example, enhance or inhibit filament growth, induce filament branching, bundling, and anchoring, or introduce motor activity into the network, a highly versatile dynamical cytoskeleton is formed.

One of the most important cytoskeletal adaptor proteins is Coronin-A. It is a single *β*-propeller-like protein linked to a coiled-coil domain [4]. It exists in all animal cells, protists and fungi, but has not been identified in plant genomes [5]. Its major function is to modify the branching and cross-linking of actin fibers by acting as a spacer while directing the Arp2/3 complex. When Coronin-A replaces Arp2/3 in older actin networks, it destabilizes the junction and provides more actin depolymerization sites [6]. It also enhances the actin severing by actin depolymerization factors [6–8].

Coronin-A was first identified in the social amoeba *Dictyostelium discoideum* (*D. discoideum*) [9]. *D. discoideum* is a single-celled motile microorganism that aggregates and differentiates upon starvation to form a multicellular fruiting structure. Due to its biochemical and morphological similarities to cancer cells and cells of the mammalian immune system, *D. discoideum* has become a widely used model for studying amoeboid motility and the dynamics of the actin cytoskeleton. Cells lacking Coronin-A show defects in protrusion formation, motility, and development [10, 11]. In *D. discoideum*, a large number of multinucleated cells was found due to insufficient mitosis in the absence of Coronin-A; in addition, motility was reduced, cells showed deformations of the nucleus and aberrant cell morphology [12]. Reduced nutrient uptake due to impaired phagocytosis, one of the major actin-dependent processes, resulted in slow growth and prolonged generation times [13].

Such functional studies of cytoskeletal proteins rely on gene knockouts. Mutations or deletions in the coding sequence prevent transcription of the functional full-length protein, leading to a so-called loss-of-function version of the protein. Phenotypic changes in the corresponding mutant cell line may then indicate the functional role of this protein. However, unexpected side effects of the remaining parts of the protein or adaptation and compensation effects by overexpression of other players in the signaling network may complicate interpretation of the observed mutant phenotype [14–17]. To avoid premature conclusions, a set of complementary mutation techniques can be applied to substantiate the results, each with its advantages and disadvantages. For example, relying on the well-established RNA interference (RNAi) technique, the level of translation can be diminished, thereby creating a knockdown mutant with a greatly reduced amount of the protein of interest. Alternatively, the knock-sideways technique (also known as chemically-induced dislocation or CID) allows for the rapid depletion of the target protein from its site of action by mislocalization to a different part of the cell. In this way, the function of essential proteins can be investigated, including immediate loss effects without giving time for compensation and adaptation effects to arise. However, in contrast to gene knockouts, possible side effects by the expression and anchoring of the dimerization system have to be considered.

This work provides a case study, in which we explore the expression and function of Coronin-A via different mutation strategies at different levels of gene or protein expression. First, we address selection-linked integration (SLI) as a genetic mutation method to obtain a stable Coronin-A knockout cell line in *D. discoideum*. For comparison, we contrast this cell line to a presumptive CorA-null cell line constructed by gene disruption, for which revertants have been described to occur previously [12]. We furthermore study a CorA-RNAi knockdown, and investigate CorA-null cells as well as *corA*-SLI cells transcribing the RNAi construct. Moreover, we use the SLI technique to establish a CID-based knock-sideways system in *D. discoideum* (CorA-KS cell line), revealing effects of a sudden loss of function of our protein of interest, expressed under its native promotor. The four cell lines (*corA*-SLI, CorA-null, CorA-RNAi and CorA-KS) are characterized and compared in terms of different actin-dependent processes: cell motility, cell morphology (shape and size), growth rates as well as their development in terms of spore and fruiting body formation. Our comparative investigation of actin-dependent processes at different scales as well as the use of complementary mutation techniques enables us to evaluate their application possibilities, benefits, strengths and weaknesses.

### Selection-linked integration for loss of function and CID

Selection-linked integration (SLI) is a mutation method that is based on the integration of an entire plasmid into the locus of a gene of interest by homologous recombination. With this approach, a loss-of-function gene knockout can be created in a fast and effective way. For a gene knockout, a homology region consisting of the first 700 bp of the gene of interest is used, which is cloned into a vector as the target sequence. This is followed by a 2A skip peptide sequence that causes an intermittent translation and allows multi-gene expression under the same promoter from a single strand of mRNA [18, 19]. The resistance gene does not include its own promoter, which provides the opportunity to select for mutated cells only [20]. Figure 1A illustrates the custom-made plasmid that was used in our study together with the gene encoding Coronin-A before and after integration of the knockout plasmid.

**Figure 1.**
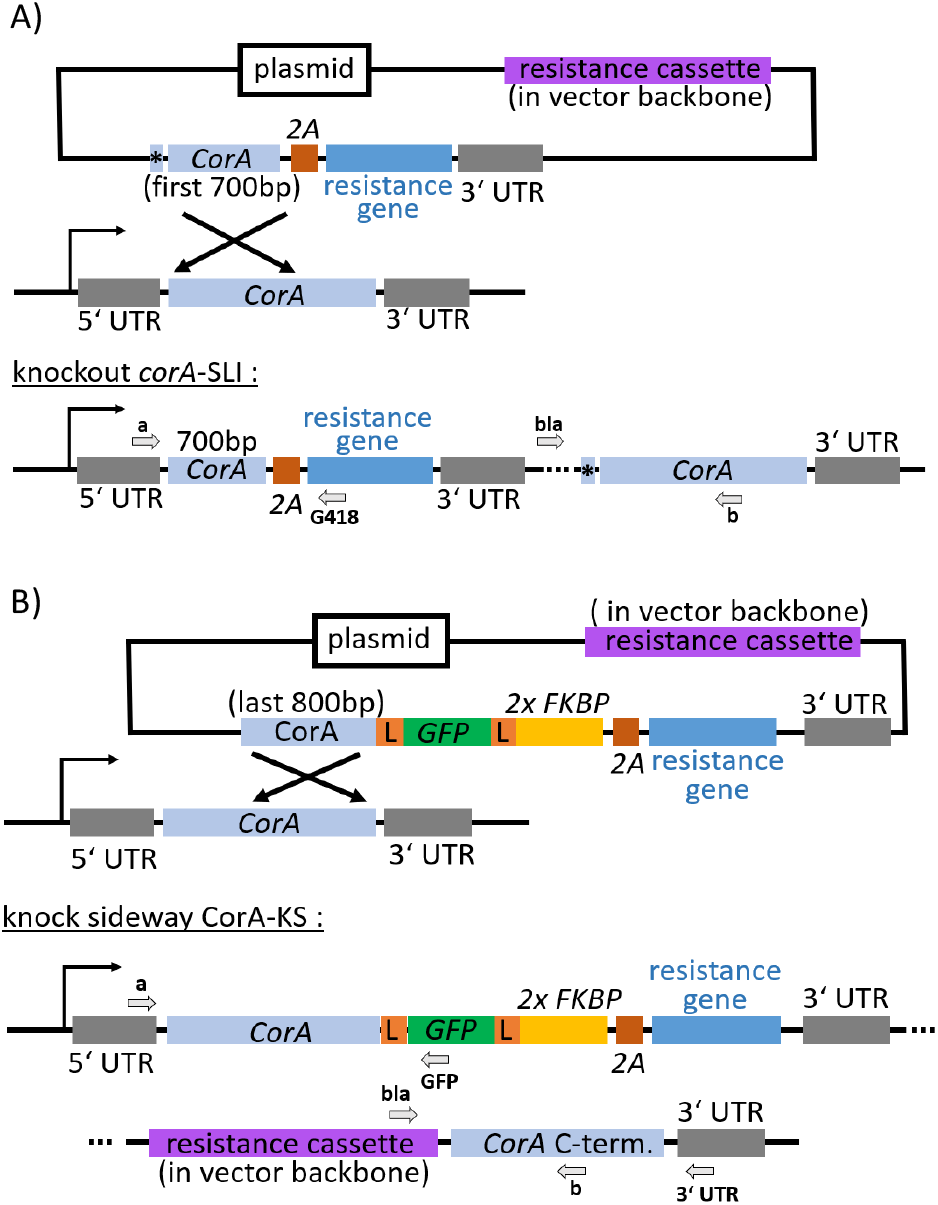
Selection-linked integration (SLI). (A) Schematic of SLI knockout strategy. (B) Schematic of SLI knock-sideways strategy. 5’ UTR, upstream untranslated region; *CorA*, gene coding for Coronin-A; 3’ UTR, downstream untranslated region; 2A, T2A skip peptide; resistance gene, Neo-R, neomycinresistance; asterisk, stop codons; GFP, green fluorescent protein coding region; L, linker; FKBP, rapamycin binding domain; arrows, promoters; filled arrows, primers: a, forward endogenous primer; b, reverse endogenous primer; bla/bla3’, forward integration check primer; G418/GFP, reverse integration check primer; *CorA* c-term., endogenous C-terminus of *CorA* gene.

Similar homologous recombination-based strategies to knock in genes are already known, and a related method has been used before in *D. discoideum* to express a tagged protein under its native promoter [21]. The SLI method that is introduced here to *D. discoideum* extends this approach so that proteins can be linked to any tag of interest, including fluorescent tags and binding domains for CID. For CID of Coronin-A, its last 800 bp without the stop codon were cloned into a knock-sideways vector that already carried coding sequences for linkers, a fluorescent protein, and the CID domains as well as a resistance gene, illustrated in Figure 1B.

## RESULTS

### SLI-based knockout differs in morphology, growth and development from CorA-null mutant and RNAi knockdown

Comparing the appearance of wild type AX2, the knockdown by RNAi, and the SLI-based knockout mutant cells, only minor differences in cell size were observed, while the gene disruption CorA-null mutant showed enlarged, multinucleated cells with 3-4 nuclei (Figure 2A, Tables S1 and S2). We performed a quantitative analysis of cell size and circularity for over 1000 cells per strain (Figure S4). Our analysis confirmed that the RNAi knockdown cell line and the SLI knockout mutant showed similar cell size distributions. They were, however, significantly different from the size distribution of wild type cells, as revealed by statistical tests. The size distribution of CorA-null mutant cells reflected the higher number of abnormally large, multinucleated cells. Consistent with these results, the RNAi knockdown and the SLI mutant also showed a similar circularity, which was larger compared to the wild type. The circularity of the CorA-null mutant cells was enhanced but did not differ significantly from the wild type.

**Figure 2.**
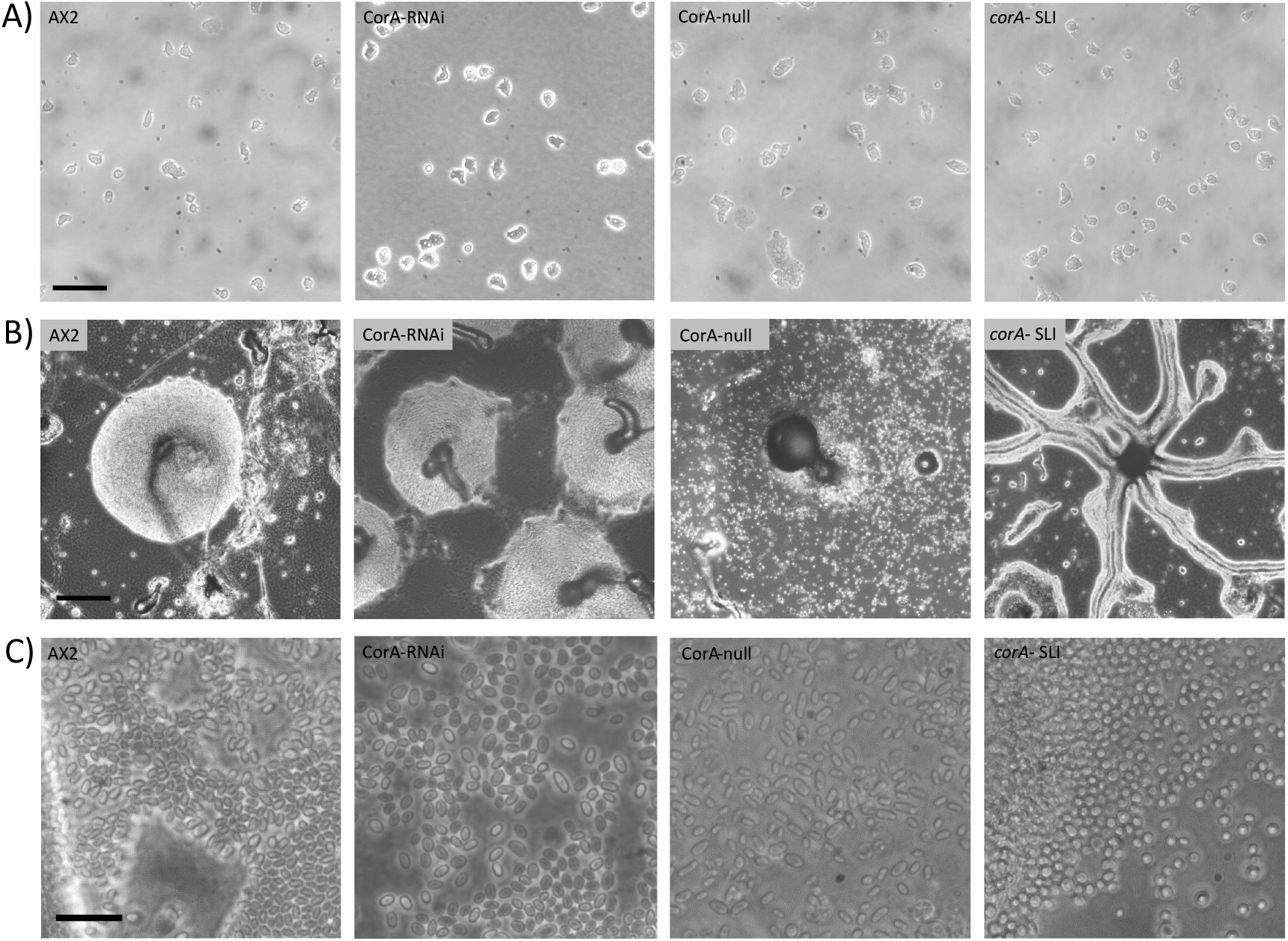
Influence of Coronin-A on cell appearance, development of fruiting bodies, and spore formation. Representative examples are displayed of AX2 wild type, CorA-RNAi knockdown, CorA-null mutant, and *corA*-SLI knockout cell lines (from left to right), showing (A) single cells for morphological analysis and determination of the cell size; (B) different stages of the aggregation process reached after 48 h; (C) appearance of spores after 48 h; scale bars: (A) and (B) 50 µm, (C) 20 µm.

The doubling time of cells increased from about 11 h in wild type to more than 20 h in both gene knockout cell lines, where Coronin-A was no longer functional (CorA-null and *corA*-SLI). In contrast, it was marginally affected by a reduced amount of the protein, as there was no decreased growth rate observed in RNAi-treated cells, see Figure S5.

Since Coronin-A is known to be upregulated during the starvation and differentiation phases of the *D. discoideum* life cycle [9, 11], we also monitored the process of fruiting body formation in all cell lines, see Figure 2B. While the AX2 wild type cells were able to differentiate and form large fruiting bodies within 24 h, the RNAi knockdown mutants showed a reduced differentiation capacity. However, they were still able to form Mexican hat-like structures within 48 h of the onset of starvation, see Figure 2B, second from left. Pronounced differences were found for the two *corA* knockout mutants: after 48 h, the CorA-null mutants formed precursors of stalk and fruiting bodies, while most *corA*-SLI cells were still in the aggregation phase.

Besides delayed differentiation, all strains retained the ability to produce spores, see Figure 2C. The CorA-null spores were of normal elongation, shape and size, similar to the AX2 wild type spores. The CorA-RNAi cell line produced smaller spores of more heterogeneous size and shape. The spores of the SLI mutant were distinctly different from spores of the other cell lines. They were smaller in size and round in shape, see Figure 2C, right. Also, the viability of the SLI mutant spores was strongly reduced compared to the other cell lines: only a few cells emerged from a large number of SLI mutant spores.

### Motility of the SLI-based knockout is reduced

In this study, we characterized the random motility of cells in terms of their mean-square displacement (MSD), quantifying how the average squared distance, which a cell has traversed with respect to its initial position, grows as a function of time. For the temporal dependence of the MSD of *corA*-SLI mutant cells and wild type, we observed curves of similar shape but the MSD of the mutant was shifted to lower values, see Figure 3A. This indicates an overall smaller effective diffusivity of the SLI mutant cells compared to the wild type. As expected, the MSD of the CorA-RNAi knockdown was found in between the curves of the wild type and the SLI knockout. For the CorA-null mutant, the MSD is close to the MSD of the SLI mutant at short time intervals and approaches the MSD of the wild type for longer times.

**Figure 3.**
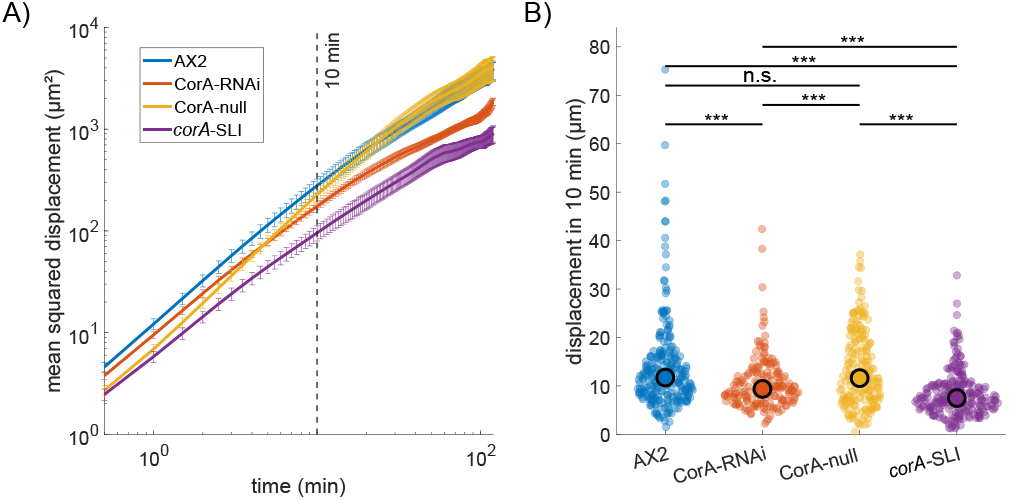
Time evolution of MSD and displacement distributions. (A) Log-log plot of MSD*±* SE versus time and (B) distributions of mean cell displacement for a time interval of 10 minutes of AX2 wild type, CorA-RNAi knockdown, CorA-null mutant, and *corA*-SLI knockout cell lines. Whereas the long-time motility of CorA-null cells equals the level of AX2 wild type, *corA*-SLI spreads significantly slower; the motility of CorA-RNAi cells lies between the levels of wild type and *corA*SLI knockout cells. Around 100 tracks were available for each strain from *n*≥ 6 experiments (for details, see Table S3). In panel (B), black circles indicate median values of the distributions. The statistical significance of differences among cell lines were assessed using Kolmogorov-Smirnov tests; *p*-values are indicated by asterisks: *** for *p* ≤ 0.001, ** for *p* ≤ 0.01, * for *p* ≤ 0.05 and not significant (n.s.).

We furthermore quantified motility in terms of the distribution of mean cell displacement within a fixed time interval. For a time lag of 10 min, a direct comparison of these displacement distributions for all four strains shows no significant difference between the wild type AX2 and the CorA-null mutant, while the *corA*-SLI displayed a significantly reduced motility (Figure 3B).

### Assessment of Coronin-A fragments in CorA-null and corA-SLI cells

CorA-null mutant cells are known to express a truncated version of the Coronin-A protein [12]. Also *corA*-SLI cells could express a nonfunctional truncated part of Coronin-A. To probe for potential side effects, we transformed an RNAi plasmid into both mutant cell lines (*corA*-SLI and CorA-null), in which the RNAi construct was directed against the N-terminal part of the*CorA* mRNA. We would thus expect the RNAi plasmid to reduce the expression of the Coronin-A fragment.

Strikingly, the *corA*-SLI mutant expressing the RNAi construct showed no significant changes in motility when compared to the *corA*-SLI mutant, but the CorA-null cells treated with RNAi closely mimicked the results for the *corA*-SLI knockout (Figure 4): the motility was similar in the *corA*-SLI mutant with and without RNAi expression, and also RNAi expression in CorA-null cells resulted in similar levels of motility with no significant differences compared to the *corA*-SLI mutant; the motility of CorA-null cells and *corA*-SLI, in contrast, is significantly different. During starvation, CorA-null cells additionally expressing the RNAi construct were no longer able to stream. They formed small and very few fruiting bodies, and the spores showed a roundish shape, as in the *corA*-SLI knockout (see Figures S7 and S8); moreover, the results of the cell circularity analysis are also similar to the *corA*-SLI mutant results (Figure S9).

**Figure 4.**
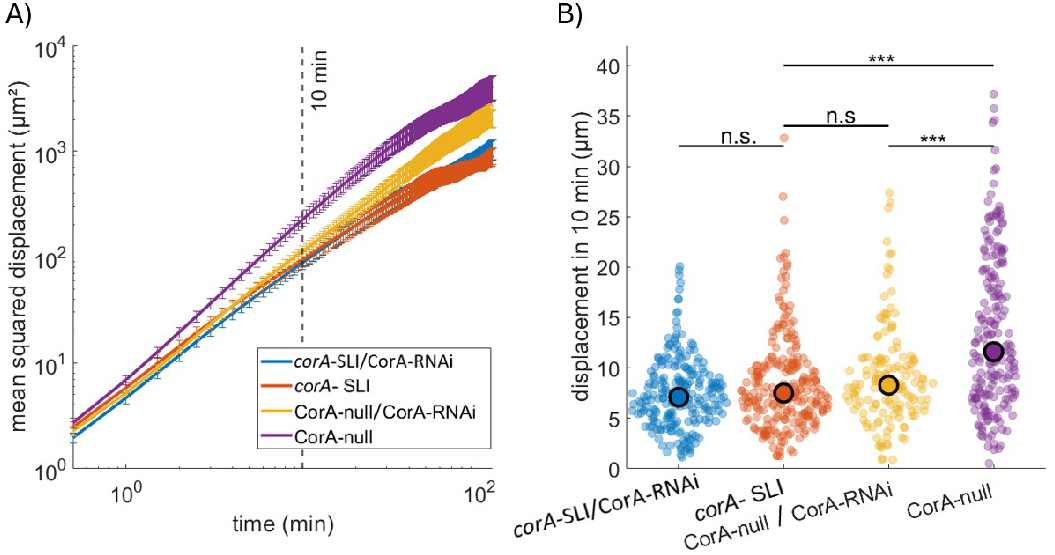
Time evolution of MSD and displacement distributions. (A) Log-log plot of MSD±SE and (B) comparison of displacement distributions after 10 minutes of *corA* gene knockout (*corA*-SLI) and the CorA-null cell line, with and without transcribing the RNAi construct. The transcription of the RNAi construct in CorA-null cells reduces the motility to the knockout level of *corA*-SLI; notably, the motility of the latter is not influenced by the RNAi construct. For technical details of the illustration, see Figure 3.

Based on these phenomenological observations, we hypothesize that the translation of the N-terminal part of the *CorA* gene is silenced in the *corA*-SLI mutant, whereas the CorA-null mutant expresses a truncated version of the protein [12], which affects the phenotypic features studied here.

### CID to induce acute loss of function

The CID mechanism is based on the reversible dimerization of two protein domains by a small linking molecule, in this case, rapamycin. To induce an acute loss of function, the protein of interest, here Coronin-A, is tagged with a fluorescent protein and an additional peptide with high affinity to rapamycin. A second protein that is expressed from another plasmid and is localized away from the site of action of Coronin-A, here at the surface of mitochondria, carries another peptide with high affinity to rapamycin to serve as an anchor for the dislocation process. When rapamycin is added, both high-affinity domains will bind to it, forming a heterodimer. As one of the binding partners is anchored to the surface of mitochondria, Coronin-A that is attached to the other binding partner, will be mislocalized to the anchoring structure and thus removed from its native site of action in the cell cortex (referred to as a “knocksideways”) [22, 23]. Since the dislocation happens at a timescale of seconds to minutes, the cell cannot adapt to the sudden loss of the protein of interest, thereby enabling a more direct assessment of its importance compared to a genetic knockout. Note, however, that side effects induced by the dimerization tags have to be considered and can influence the results of CID studies.

We explored the SLI method to design a CID system for the mislocalization of Coronin-A. The approach to design the knock-sideways system was similar to the SLI knockout construction, but here the last 800 bp of the *CorA* gene served as the homology region. The 3’ end of the gene in the plasmid lacked the stop codon but was linked to the green fluorescent protein (GFP) coding sequence and to one of the dimerization domains. We used 2x FKBP12 (linkage of two times FK506 binding domain), which is known to bind to rapamycin [22]. The FKBP12-rapamycin complex binds to the FRB (FKBP and rapamycin binding) domain of TOR kinase linked to a protein of the outer mitochondrial membrane [24]. Thus, when rapamycin was added to the cell culture, the tagged Coronin-A translocated to the mitochondria and was no longer available at its native site of action, resulting in a sudden loss of function.

A schematic of the knock-sideways system is displayed in Figure 5A, with the inactive CID system on the left and the rapamycin-activated system on the right. In Figure 5B, CorA-KS cells expressing the knock-sideways constructs containing GFP-tagged Coronin-A and a red actin marker (LimE-RFP) are shown with and without activated CID. Co-localization of the tagged actin polymerization marker LimE with Coronin-A is observed in the top row. The activated CID system 22 min after addition of a final concentration of 10 µM rapamycin is shown in the bottom row. Most of the cortically localized Coronin-A was bound to mitochondria (green dots), while some cortical actin can still be detected at pseudopodia based on the LimE-reporter (Figure 5B).

**Figure 5.**
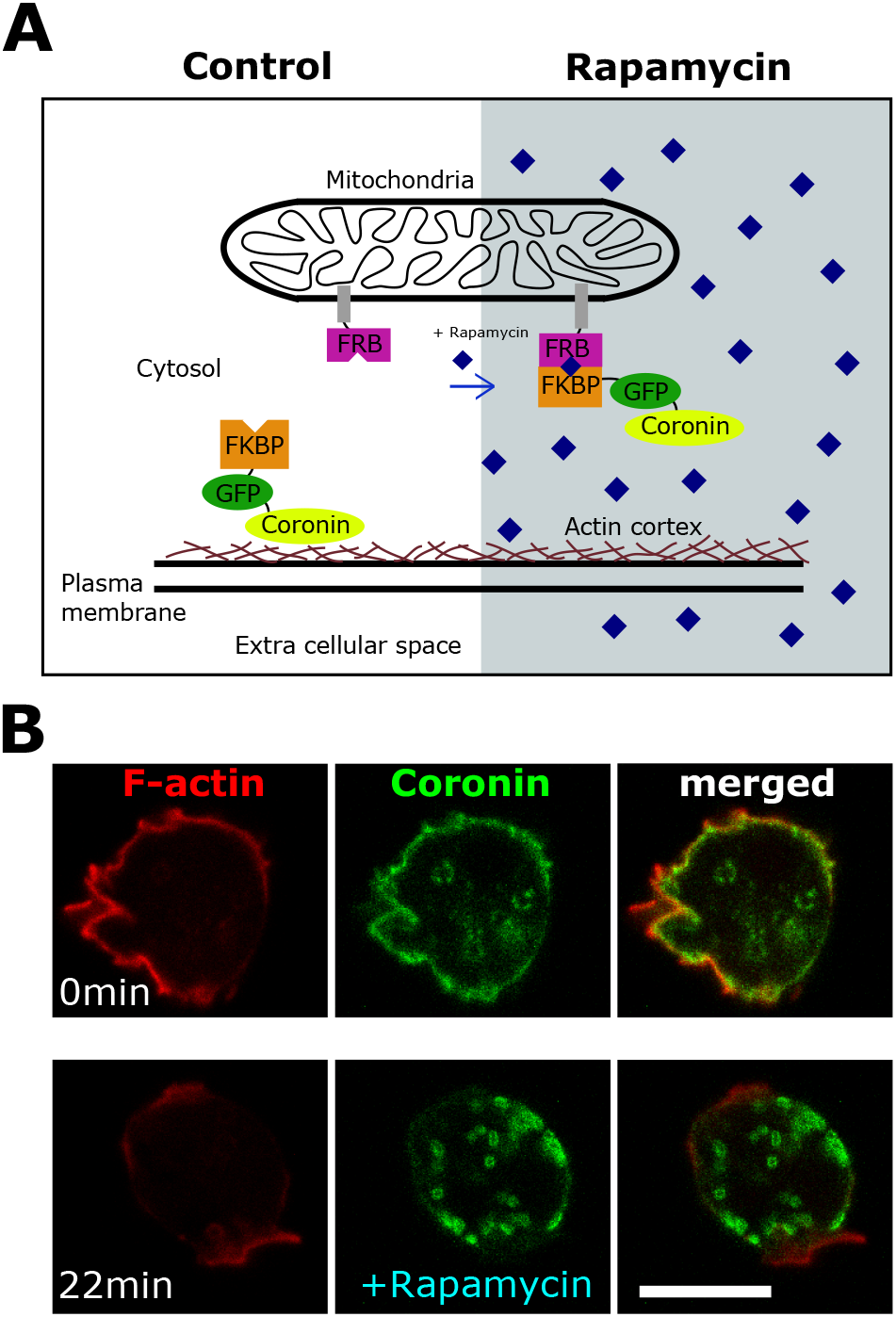
Knock-sideways of Coronin-A. (A) Schematic of the knock-sideways system in *D. discoideum*. The FRB domain localized to the mitochondria via an N-terminally fused signal peptide. The Coronin-A was tagged C-terminally with a GFP and a FKBP domain, and colocalized with F-actin. Upon addition of rapamycin, FKBP and FRB domains dimerized and Coronin-A was trapped at the mitochondria and could not interact with F-actin at the cortex. (B) Representative *D. discoideum* cell expressing LimE-RFP (red channel, F-actin), Mito-FRB, and Coronin-GFP-2xFKBP (green channel). At the beginning of the experiment (0min), rapamycin (10 µM) was added. Before addition of rapamycin, Coronin-A colocalized with F-actin at the cell cortex. 22 min after rapamycin addition, Coronin-A was predominantly found at the mitochondria. Scale bar is 10 µm.

### Expression of CID system does not affect streaming and fruiting body formation

The correct integration of the SLI constructs into the *D. discoideum* host genome was confirmed by polymerase chain reaction (PCR) based on specific primer pairs (Figure S3). The full-length expression of the knock-sideways Coronin-A and a complete substitution of the endogenous Coronin-A was validated by immunoblotting with an anti-Coronin-A antibody that binds to the C-terminal part of the protein (Figure S2). The expected molecular weight of native Coronin-A of 49 kDa and of the tagged knock-sideways version consisting of Coronin-A, GFP, and 2x FKBP domains of approximately 90 kDa were identified by Western blotting (Figures S2). The amount of the knock-sideways construct equalled the amount of Coronin-A expressed under its endogenous promoter, as indicated by comparable strengths of the two bands (Figures S2).

Cells expressing the CID-based knock-sideways system (CorA-KS mutant cells) were not impaired or delayed in streaming and fruiting body formation while the CID system was inactive, although their spore-containing heads were smaller than the wild type (Figure S6). The shape and viability of the CorA-KS mutant spores were similar to those of the CorA-RNAi mutant. They showed a more heterogeneous phenotype than the wild type, but were not as rounded as the knockout mutant spores (Figure 2C and Figure S6). Growth and division under vegetative conditions proceeded much slower compared to the wild type (Figure S5). We measured the cell densities in liquid HL-5 shaking culture over a period of 48 h and calculated the doubling times. For AX2 wild type cells, we measured a doubling time of 11.3 h (Figure S5), which was larger than values of around 8 h, reported previously [25]. However, with a culture temperature of 20 °C that was used in our experiments and given the impact of media composition and selection drugs [26], this value still matches the published range from 8 h at 22 °C to 12 h at 19 °C [25, 26]. Expression of the knock-sideways constructs with tagged Coronin-A resulted in a pronounced increase of the doubling time to 30.3 h. In comparison, the doubling time of the *corA*-SLI knockout was 20.9 h (about twice as long as for the wild type), while the RNAi-treated cell line did not show an increase in doubling time at all (Figure S5). Thus, low levels of Coronin-A were sufficient to maintain growth rates and development, while expression of the knock-sideways construct severely impacts replication times.

### CID of Coronin-A reduces motility to the knockout level

We compared the motility of the Coronin-A knocksideways cells in the presence and absence of rapamycin with the AX2 wild type and the SLI-based gene knockout of Coronin-A. As rapamycin may influence motility independent of its function as a mediator of CID [27], also the motility of AX2 wild type and SLI knockout cells was measured both in the presence and absence of rapamycin. To quantify changes in motility, we again relied on the time evolution of the MSD and the displacement distributions for a time lag of 10 minutes. When comparing the MSD curves of the investigated cell lines, we found that treatment with a final concentration of 10 µM rapamycin shifts the MSD curves of both AX2 wild type and SLI knockout cells to higher values compared to cells of the same strain in the absence of rapamycin, indicating that rapamycin enhances motility, see Figure 6A. The two MSD curves of rapamycin treated and untreated cells are only similar in the case of the Coronin-A knock-sideways cell line (CorA-KS), suggesting that the rapamycin induced enhancement is balanced by a reduced motility due to the CID-induced acute loss of Coronin-A.

**Figure 6.**
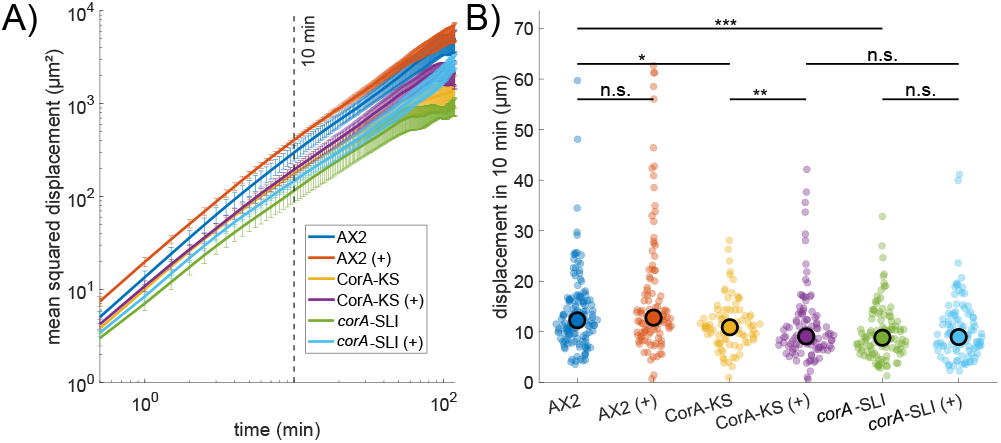
Time evolution of MSD and displacement distributions including CID-based acute loss of function. (A) Log-log plot of MSD*±* SE and (B) comparison of displacement distributions after 10 minutes of AX2 wild type (AX2), Coronin-A knock-sideways (CorA-KS), and *corA* gene knockout (*corA*SLI) cell lines, each in the presence (+) and absence of 10 µM rapamycin. Rapamycin-treated CorA-KS cells display a motility level comparable to the *corA*-SLI knockout. For technical details of the illustration, see Figure 3.

From the distributions of displacements for a time lag of 10 min (Figure 6B), the stimulating effect of rapamycin was less obvious—the median of displacement distributions is a reliable measure for motility in a heterogeneous cell population which is robust against outliers. According to the Kolmogorov-Smirnov tests, the displacement distributions of AX2 wild type and SLI knockout cells showed no significant changes upon treatment with rapamycin. Based on the displacement distribution, a significant decrease in motility was detected for the CorA-KS cells when treated with rapamycin, resulting in the rapamycin-induced sudden mislocalization of Coronin-A to mitochondria. This can be assigned to the rapamycin-induced sudden mislocalization of Coronin-A to mitochondria. Prior to rapamycin treatment, CorA-KS cells showed a small but significant difference to the AX2 wild type, which can be most likely attributed to side effects of expressing the additional dimerization tags. After rapamycin treatment, however, motility of the CorA-KS cells dropped to the level of the SLI gene knockout with no significant difference between the respective displacement distributions.

## DISCUSSION

### SLI-based knockout of Coronin-A in AX2 background

In this study, we introduce selection-linked integration (SLI) to knock out Coronin-A in *D. discoideum* as an example of an important cytoskeletal adaptor protein. We observed pronounced differences in terms of cell motility, cell morphology, and development between the previously published CorA-null mutant [12] and our SLI knockout *corA* cells. Based on our phenomenological observations, we hypothesize that these differences may be attributed to potential side effects of the truncated 200 amino acid long version of the protein, the RNA of which is expressed in the CorA-null mutant shown by northern blot analysis [12]. This assumption is based on the result that, remarkably, the motility of the SLI knockout mutant did not change when expressing the RNAi construct in comparison to the *corA*-SLI mutant, but the CorA-null cells treated with RNAi closely mimicked the results for the corA-SLI knockout. Another axenic strain, called DH1-10, has been used in the past to investigate the function of Coronin-A, however, it is unclear whether the DH1-10 *corA* knockout mutant expresses a truncated version of CorA [11]—all tested antibodies for immunoblot in this study were only directed against the C-terminal part of the protein [11]. Notably, uncontrolled reversions of the *corA* gene could occur in the CorA-null [12] or DH1-10 cells [11], but these are unlikely to happen in the SLI knockout.

Compared to the SLI knockout, the CorA-null mutant was still able to produce fruiting bodies, see Figures S7-S8, which is not the case for CorA-deficient DH1-10 cells [11]. The wild type appearance of the CorA-null spores (Figure S8) suggests that the partially expressed N-terminal Coronin-A fragment is involved in the differentiation process and the formation of fruiting bodies and spores. In addition, cells with a reverted *corA* gene could have compensated for deficiencies of mutant cells in stalk and fruiting body formation. The different shapes of spores and the large differences in viability, dormancy, and germination of spores are generally influenced by actin rods [28]. Considering that Coronin-A is involved in actin branching, we speculate that in mutants lacking Coronin-A, the hexagonal arrangement of the actin tubules, which is important for spore formation, may be inhibited [28]. This may explain the low viability of SLI mutant spores. In addition, the cell size distribution of the CorA-null mutant showed a wider spread, and aberrant mitosis resulted in large multinucleated cells and cell aggregates, which were not found in the SLI mutant cultures.

Interestingly, motility of the CorA-null cells showed a mixed behavior as revealed by the time dependence of the MSD. For lag times around one minute, the MSD of CorA-null cells resembled the MSD of the SLI knockout mutant, whereas for longer time intervals, it approached the MSD of wild type cells. This is in agreement with earlier results, where the speed of CorA-null cells, calculated from displacements over 30 s, showed a clear difference to the speed of wild type cells [12, 29]. We conjecture that the formation of *de novo* pseudopodia is affected by a loss of Coronin-A function, resulting in decreased displacements on short time scales. Cell polarity and persistence, on the other hand, could have been maintained, thus enhancing long-time displacements to the wild-type level. A possible explanation could be that this is related to the ability of Coronin-A to inhibit Cofilin, which will be affected in its truncated version due to the missing C-terminal coiledcoil domain [30]. As the coiled-coil domain is important for multimerization [31–35], loss of 2/3 of the propeller structure can also have other additional effects: it could still bind to actin but cannot interact with other proteins nor dimerize due to the missing C-terminus and will thus be blocking binding sites of other proteins.

### CID-based knock-sideways complements gene knockout to study acute loss of Coronin-A function

Knocksideways assays have already been successfully established in *D. discoideum* [24, 36, 37]. *D. discoideum* cells were reported to switch between different migratory modes when intracellular PIP_2_ and Ras levels are altered by a rapamycin-based mislocalization system [38]. Sequestration of the G protein subunit G*β* using the knock-sideways technique can tune the coupling between cytoskeletal actin oscillators [15].

A novelty of this study is the use of the SLI-technique to introduce a CID-system into *D. discoideum* to study the sudden loss of function of Coronin-A. As shown in Figure 5B, tagged Coronin-A is localized at the cell cortex if the knock-sideways system is not active, and only a small amount of the protein is bound to subcellular structures and organelles, which is caused by endogenous binding partners of the CID protein domains. Already 22 min after treatment with the dimerizing agent rapamycin, the knock-sideways system was fully active and a strong recruitment of Coronin-A to the mitochondria could be observed.

Based on this knock-sideways system, a decrease in *D. discoideum* motility could be induced. Upon addition of rapamycin, motility was reduced from wild-type level down to a level comparable to the motility of Coronin-A-deficient SLI knockout cells (note that expression of the knock-sideways tags slightly decreased motility, even without activation by rapamycin; cf. Figure 6B). We furthermore observed that rapamycin itself induced an increase in the motility of wild type and SLI knockout cells [27]. This suggested that the decrease in motility, which was induced by the sudden loss of Coronin-A function, was partly compensated by the rapamycin induced enhancement of motility.

Tagging Coronin-A with the knock-sideways dimerization domain did not influence cell differentiation and spore formation. As upregulation and dimerization of Coronin-A were reported to occur mainly in the starvation phase and appeared to be critical for the ability to stream and survive [39, 40], we conclude that a modified C-terminus of the protein does not affect these processes. Also in *Xenopus*, dimerization of Coronin was shown to be essential for its proper function and localization [35].

In contrast to the negligible effects on cell differentiation and spore formation, growth rates were strongly reduced in cells expressing the Coronin-A knock-sideways construct, even in comparison to the SLI gene knockout. This suggests that the C-terminal coiled-coil domain of the protein plays an important role in the interaction with other proteins during the growth and proliferation phase, where the knock-sideways domains could induce steric hindrance.

In summary, we have compared methods to modulate protein function by interfering at the gene, mRNA, or protein level. Relying on Coronin-A as a case study, we have demonstrated the advantages of the SLI technique focusing on the loss of Coronin-A function in an SLI-based gene knockout, and in a CID-based knock-sideways construct expressed under the native promotor of the protein of interest. From our findings we conclude that the SLI method is a suitable choice to study the impact of a Coronin-A knockout at the genetic level in *D. discoideum*.

## METHODS

### Cell culture and media

All *D. discoideum* cells were derived from the AX2-214 strain and were cultured axenically at 20 °C in untreated cell culture flasks with an area of 75 cm^2^ or in 25 ml shaking culture at 180 rpm in HL-5 media including glucose supplemented with vitamins and micro-elements (Formedium, UK) with the appropriate selection markers. Cells were pelleted, washed twice in Sørensen buffer for preservation and long-term storage, and plated on Sørensen agar plates containing 1.2% agar.

### Strains and plasmids

All plasmids were constructed using restriction enzyme digestion of double-stranded DNA and homologous recombination-based Gibson Assembly. The plasmid backbones of the two small expression vectors pDM326 for SLI constructs or pDM358 for cytosolic expression were used [41]. Both carry an actin-15 promoter and an actin-8 3’UTR. The correct sequence of all constructs was confirmed by Sanger sequencing which was performed commercially at LGC Genomics (Berlin, Germany). Transformation of the plasmid into *D. discoideum* cells was achieved by electroporation and transformants were selected with appropriate markers (Blasticidin S (hydrochloride), Cayman Chemical (AnnArbor, USA), 10 µg/ml; G418-Disulfat (Geneticin), Amresco (Solon, USA), 10 µg/ml; Hygromycin B Gold, InvivoGen (San Diego, USA), 50 µg/ml; Penicillin/Streptomycin 10X, Biological Industries (Beit HaEmek, Israel), Penicillin G SodiumSalt 100 U/ml, Streptomycin Sulfate 100 µg/ml). RNAi plasmid construction was based on the strategy of H. Martens [42]. The plasmid backbone of pIS193 was used in this work, which was kindly provided by Ralph Gräf and Irene Meyer (Department of Cell Biology, University of Potsdam, Germany; see also Ref. [43]). One part of its mCherry coding sequence between the NheI and AflII restriction sites in the plasmid was replaced by 500 bp of the antisense sequence of the 5’ part of the CorA coding gene. In the second step, the sense fragment of the same gene section was inserted between the SalI and BamHI restriction sites. 190 bp of the mCherry coding sequence was left in between to act as a linker, allowing the sense and antisense strands to bind and form a loop. Finally, the backbone resistance gene was changed from geneticin to hygromycin. The Tth111I restriction site was followed by the more commonly used ApaI restriction site. At the 3’ end of the resistance gene, an EcoRI restriction site was added in 5’ position to the BstBI restriction site.

### Growth curves

Cells were grown in a shaking culture overnight and diluted to a density of 10^5^ cells/ml. The number of cells/ml was manually counted every 8 h for 2 d using a Neubauer counting chamber. Each strain was tested in duplicate with (+) and without rapamycin at a final concentration of 10 µM. Generation times were determined by an exponential fit to cell concentrations for each experiment and subsequent averaging of the resulting growth rates.

### Streaming and fruiting body formation

After harvesting cells from a culture flask, the cell suspension was centrifuged at 500*g* for 5 min and washed twice in Sørensen buffer. The pellet was resuspended in 200 µl buffer and plated on Sørensen agar plates, containing 1.2% agar. Images were taken at 0, 24, 48, and 72 h using an Olympus CKX41 inverted microscope (LCAch N 40x/0.55na php or CAch N 10x/0.25na php objective) and a Thorlabs DCC1645C camera. The viability of the spores was not quantified, but visually inspected and estimated in comparison to the wild type.

### Motility assay and analysis

Cells were harvested and seeded in a 24-well glass bottom plate (1.9 cm^2^/well) at a density of 1.5 10^4^ cells/well. Each line was tested with DMSO as solvent mock-treated control and with rapamycin dissolved in DMSO to a final concentration of 10 µM rapamycin. Cells were allowed to settle for 20 min for induction of the drug effect. Images were captured every 30 s by using confocal laser scanning microscopy (Zeiss LSM780, Plan-Apochromat 20x/0.8 M27 air objective, with 561 nm laser at 1% intensity and ZEN software). The transmitted light detector (TPMT channel) was used to generate brightfield images, which were later segmented and cells tracked using a MATLAB^®^ (2019b, MathWorks, Natick, MA, USA) program (see below). Experiments were performed on at least six different days and pooled.

Images acquired with a confocal LSM780 (Zeiss, Germany) were exported from ZEN software in tagged image file format and cells were segmented by using MATLAB^®^ (2019b, MathWorks, Natick, MA, USA) object edge detection with the Sobel method (MATLAB routine: edge with option ‘sobel’). An order-statistical filter was applied (MATLAB routine: ordfilt2), replacing each pixel with the 8th value of its sorted 3x3 neighborhood and the resulting image was smoothed with a Gaussian filter (MATLAB routine: imgaussfilt with a standard deviation of 1.8 pixels). After normalization, the images were binarized using Otsu’s method (MATLAB routine: imbinarize with option ‘global’), holes were filled (MATLAB routine: imfill with option ‘holes’), and the objects were eroded (MATLAB routine: imerode using a one-pixel radius disk as the morphological structuring element) to obtain a correct representation of the cells. The centroid of each cell area was determined and the position of each cell was tracked based on [44] from frame to frame over the entire image sequence of 240 frames (2 h, 30 s time interval). Finally, approximately 100 tracks per strain (see Supplementary Table S3 for detailed numbers) were saved and plotted as an overlay on the brightfield images. In a final step, the numbered tracks were manually checked for floating or dead cells in each measurement. These tracks were then removed from the track list.

For each trajectory **r**_*j*_ (*t*) with a track index *j* and the time *t*, the time-averaged mean squared displacement (TAMSD) was calculated as follows:

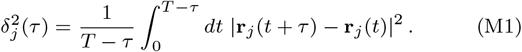

Afterwards, these values 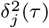 are averaged using the arithmetic mean over all cells (track indices *j*) measured on the same day (ensemble-average of TAMSD, denoted as ⟨TAMSD⟩, as also described in Refs. [45, 46]. The mean of ⟨TAMSD⟩ of different days was then calculated and plotted in a log-log plot. The standard error was determined and plotted to give an indication of the reproducibility of the results between different days.

In addition, for each cell *j* the mean of the displacements | **r**_*j*_ (*t* + *τ*) − **r**_*j*_ (*t*) | within a given lag time *τ* was calculated and illustrated in a violin plot showing a histogram of these values. The prominent larger dots indicate the median of the distribution, which is statistically more robust to outlying values than the mean.

### Morphology and cell size

To examine the morphology and cell size of all strains, adherent cells were harvested and seeded with a density of 2 10^4^ cells/cm^2^. After 4 h brightfield microscopy pictures were taken using an Olympus CKX41 inverted microscope with a LCAch N 40x/0.55na php objective and a Thorlabs DCC1645C camera. A minimum of 1000 cell outlines per strain were manually marked in red in these images using an Apple Pencil^™^(2. Gen) on an Apple iPad Pro (11”)^™^. The manually marked cell outlines in the brightfield images of at least 1000 cells were processed in MATLAB^®^ (2019b, MathWorks, Natick, MA, USA) 2019b. To do this, the manually labeled cell contours were read from the red channel of the labeled images, the resulting images were binarized (MATLAB routine: imbinarize), and the resulting contours were filled using a morphological filling method (MATLAB routine: imfill with the ‘holes’ option). The binarized image was then segmented (MATLAB routine: bwconncomp identifying connected components). For each coherent set of pixels, the centroid, area, and circularity were calculated (MATLAB routine: regionsprops with options ‘centroid’, ‘area’, and ‘circularity’). The number of connected pixels of the objects was converted to cell size according to the magnification factor used. Circularity *C* is defined as

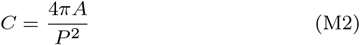

with the area *A* and the perimeter *P*, such that for a perfect circle, where *A* = *πr*^2^ and *P* = 2*πr*, the circularity is *C* = 1. For all other shapes *C <* 1.

### Confocal LSM image acquisition with Airyscan

To verify the expression of the desired rescue construct of modified Coronin-A and LimE in mutant *D. discoideum* cells, fluorescence images of transformed cell lines were captured using the ZEN software on a ZEISS confocal LSM 880 with Airyscan mode and Plan-Apochromat 63x/1.4 Oil DIC M27 objective to obtain better quality images. The 488 nm laser line was used with 4-6% laser power for GFP excitation and the 561 nm laser with 4% laser power for mRFPmars excitation to visualize GFP-tagged Coronin-A and mRFPmars-LimE in knock-sideways lines.

### Proteinextraction and immunoblot

Two million cells per strain were resuspended in 50 µl lysis buffer (9 M Urea, 10% (w/v) SDS, 5% (v/v) *β*-Mercaptoethanol, 0,2% (w/v) Bromophenol blue) and incubated for 5 min at 70 °C. The derived crude protein extract was stored at -20 °C. Discontinuous SDS-PAGE was performed using a Biorad mini-gel system chamber. 12.5% polyacrylamide gel with a depth of 1 mm was run for 1.5 h at 120 V. A molecular weight standard (Thermo Scientific; PageRuler Prestained Protein Ladder; #26616) was run together with 25 µl of each sample. After the run, the transfer was performed either overnight at 15 V or for h at 100 V in an ice bath. The transfer was performed in Towbin buffer with a nitrocellulose membrane (Amersham Protran 0.45, GE Healthcare Europe GmbH, Freiburg, Germany). The blot was blocked with 5% (w/v) milk powder in TBST and additionally with 5% (w/v) BSA in TBST. Primary antibody staining was performed overnight at 4 °C or for 1-2 h at room temperature. All antibodies were diluted in 5% (w/v) BSA in TBST according to their recommended concentration (anti-Coronin, AB 10571321, 176-3-6, 26 µg/ml, DSHB (Iowa City, USA), 1:5000; anti-Actin, AS13 2640, Agrisera (Vännäs, Sweden), 1:3000). After removal of the unbound primary antibody, the appropriate HRP-conjugated secondary antibody (goat-anti-rabbit, AP307T, Merck (Darmstadt, Germany), 1:13000; goat-anti-mouse, AS10 1189, Agrisera (Vännäs, Sweden), 1:10000; streptavidin-HRP conjugate, RPN1231, Merck (Darmstadt, Germany), 1:5000, Streptavidin-HRP conjugate was used to detect the highly biotinylated mitochondrial marker protein MCCC1 (3-methylcrotonyl-CoA carboxylase *α*) as an additional loading control to actin [47], which could be affected by varying levels of Coronin-A) was diluted in 5% (w/v) BSA in TBST and the membrane was incubated for 1-2 h at room temperature. After removing the unbound secondary antibody and Streptavidin-HRP conjugate, the substrate solution was added to the blot, and the emitted light was detected using a Fujifilm Intelligent Dark Box employing an Atik 460 EX camera.

## ACKNOWLEDGMENTS

This research has been partially funded by Deutsche Forschungsgemeinschaft (DFG), grant 318763901—SFB1294 (RG and CB). We gratefully thank Kirsten Sachse for her technical work in the laboratory. We particularly thank Ralph Gräf and Irene Meyer for providing the RNAi plasmid and many productive discussions.

## AUTHOR CONTRIBUTIONS

MS did all experiments, analyses and wrote the paper, RG programmed all analysis tools and wrote the paper, SF introduced the SLI technique, MG and CB supervised the work and wrote the paper.

## Supplementary Information

Accompanies this paper.

### Competing financial interests

The authors declare no competing financial interests.

## SUPPLEMENTAL MATERIAL

### Impact of rapamycin on cell size and shape is reduced in Coronin-A-deficient cells

When grown in the presence of rapamycin, the size and shape of *D. discoideum* cells were affected. AX2 wild type cells responded to rapamycin treatment by forming larger cells on average, as shown by the size distribution functions (Figure S1A). The circularity of wild type cells was also increased in the presence of rapamycin and reached the level of the SLI knockout cells (Figure S1B). For the CID knock-sideways and the SLI knockout cell lines, no significant differences in the cell size distributions were observed between growth in the presence and absence of rapamycin. Cells of the SLI knockout were, however, larger on average than the CID knock-sideways cells (Figure S1A). The shift in circularity was also more pronounced in the knock-sideways cell line, while the circularity did not change much in the SLI knockout under rapamycin treatment (Figure S1B). These observations confirmed the role of Coronin-A in maintaining an intact actin cytoskeleton, so that cells appeared larger and more rounded if Coronin-A function was impaired. Overall, the side effects of rapamycin with respect to cell size and shape were more pronounced in the wild type than in Coronin-A knockout cells, where cytoskeletal integrity was already compromised.

### Limits of CID

Can be found for very strong phenotypes e.g. in double mutants. The Aip1-null#9.1 cell line [1] was used as mutant background to generate an *aip1/corA* double knockout and an *aip1* /CorA-KS double knock-sideways by the SLI method. The growth and development of double knockouts is highly impaired, as expected from previous studies on comparable double knockouts [2]. The generation time is 10 h slower than the one of the Aip-null#9.1 cells alone. Double mutant cells were much larger in size due to aberrant or incomplete mitosis. Cells of the double knockout were unable to stream under starvation conditions, resulting in a complete loss of the ability to produce spores. To avoid working with this cell line, the Coronin-A knock-sideways was introduced to the Aip1-null#9.1 cells. The generation time of the double knock-sideways cells with inactive CID system was comparable to the Aip1-null#9.1 single mutant cells, which was an advantage over the double knockout. Under rapamycin treatment, the knock-sideways double mutant cells hardly proliferated at all. Whether this is an indication of the stronger phenotype in the non-compensating cell lines is not clear. In comparison to the double knockout, the double knock-sideways cells are able to stream and produce fully functional fruiting bodies with viable spores comparable to the Aip-null#9.1 mutant. The cell size distribution is indistinguishable from the wild type and single Aip-null#9.1 mutant, but the untreated cells show a more rounded shape similar to the double knockout cells. Besides all these advantages in cell culture and proliferation, the motility of the untreated knock-sideways cell line and the double knockout cell line is similarly reduced compared to the Aip-null#9.1 mutant cell line.

### Estimation of Coronin-A expression in CorA-RNAi cells based on Western blots

Figure S2 represents an exemplary Western blot, used to test the expression of Coronin-A in all cell lines of interest. Whereas no native Coronin-A was detected for *corA*-SLI, CorA-KS and CorA-null mutant cells, there still is a significant band for the CorA-RNAi knockdown. However, the band size of loading controls differ as well among the cell lines. To estimate the reduction of Coronin-A expression in CorA-RNAi cells in comparison to the wild type level, we compare the size of bands with respect to the mitochondrial marker protein MCCC1 (3-methylcrotonyl-CoA carboxylase *α*) which is used as a loading control. Note in this context that actin cannot serve as a loading control for quantitative analysis due to potential interferences with the concentration level of Coronin-A.

The Coronin-A concentration is assumed to be proportional to the size of bands, which was determined by selecting a region of interest of fixed size around each band first, followed by background subtraction and summation of the pixel intensity. In three tested cases, we estimated an approximate reduction of Coronin-A concentration to 62 %, 79 % and 72 %, respectively, implying an average Coronin-A concentration level of 71 % in CorA-RNAi cells with respect to the wild type. We hence conclude that there is a significant reduction of Coronin-A in knockdown cell lines, while complete silencing could not be achieved.

**Figure S1.**
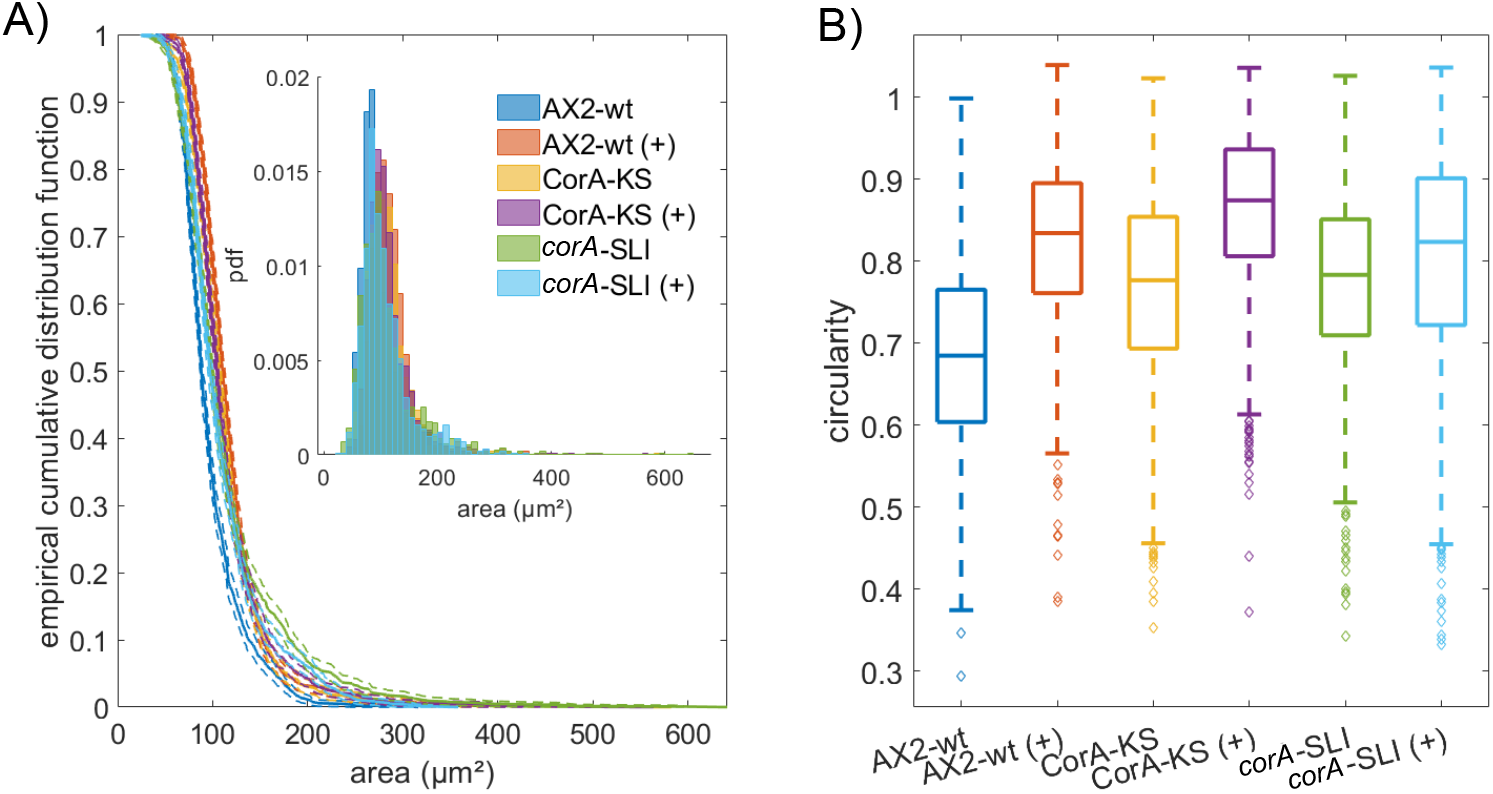
Impact of rapamycin on cell size and circularity. (A) Cell size distributions and (B) cell circularity of AX2 wild type (AX2), Coronin-A knock-sideways (CorA-KS), and *corA* gene knockout (*corA*-SLI) cells after 72 h of growth in the presence (+) or absence of 10 µM rapamycin. A minimum of 1000 cell shapes and sizes were examined per strain. For statistical analysis results of Kolmogorov-Smirnov test, see Tables S1-S2.

**Figure S2.**
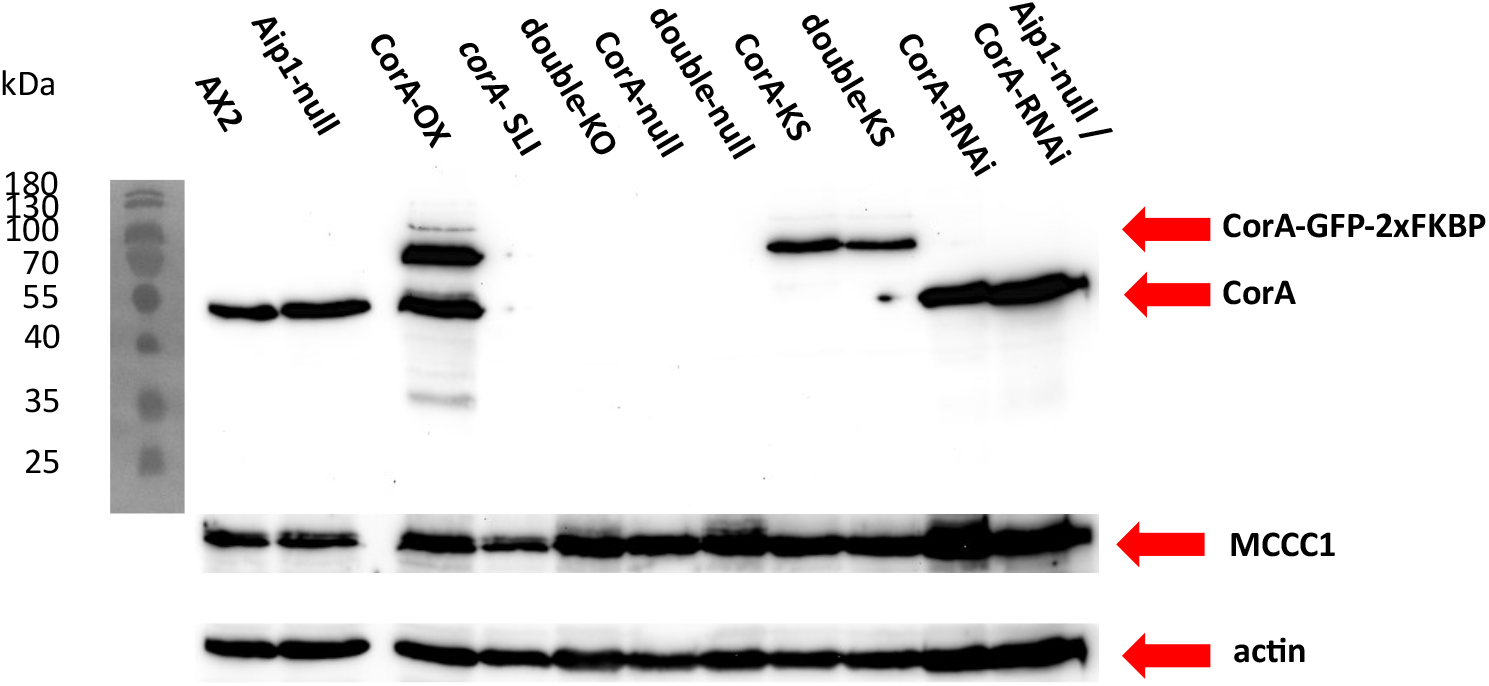
Western blot (immunoblot) for detection of Coronin-A. On lysates of wild type AX2 cells, *corA* single and Aip1null/*corA-* SLI double knockouts and knock-sideways as well as loss-of-function cell lines, CorA overexpressor and RNAi in AX2 and Aip1-null background; wild type AX2, Aip1-null#9.1 mutant [1]; CorA-OX: knock-sideways overexpressor; *corA*SLI: *corA* knockout done with SLI method; double-KO: *corA* knockout done with SLI method in Aip1-null mutant; CorA-null: classical gene disruption mutant of *corA* CorA-null HG1569#1 [3]; double-null: Aip1-/CorA-strain [2]; CorA-KS: Coronin-A knock-sideways done with SLI method; double-KS: Coronin-A knock-sideways done with SLI method in Aip1 mutant; CorA-RNAi: knockdown; Aip1-null/CorA-RNAi: knockdown of CorA in Aip1-null background; MCCC1 and Actin as loading controls. The Western Blot displays one representative out of three experiments with highly similar results.

**Figure S3.**
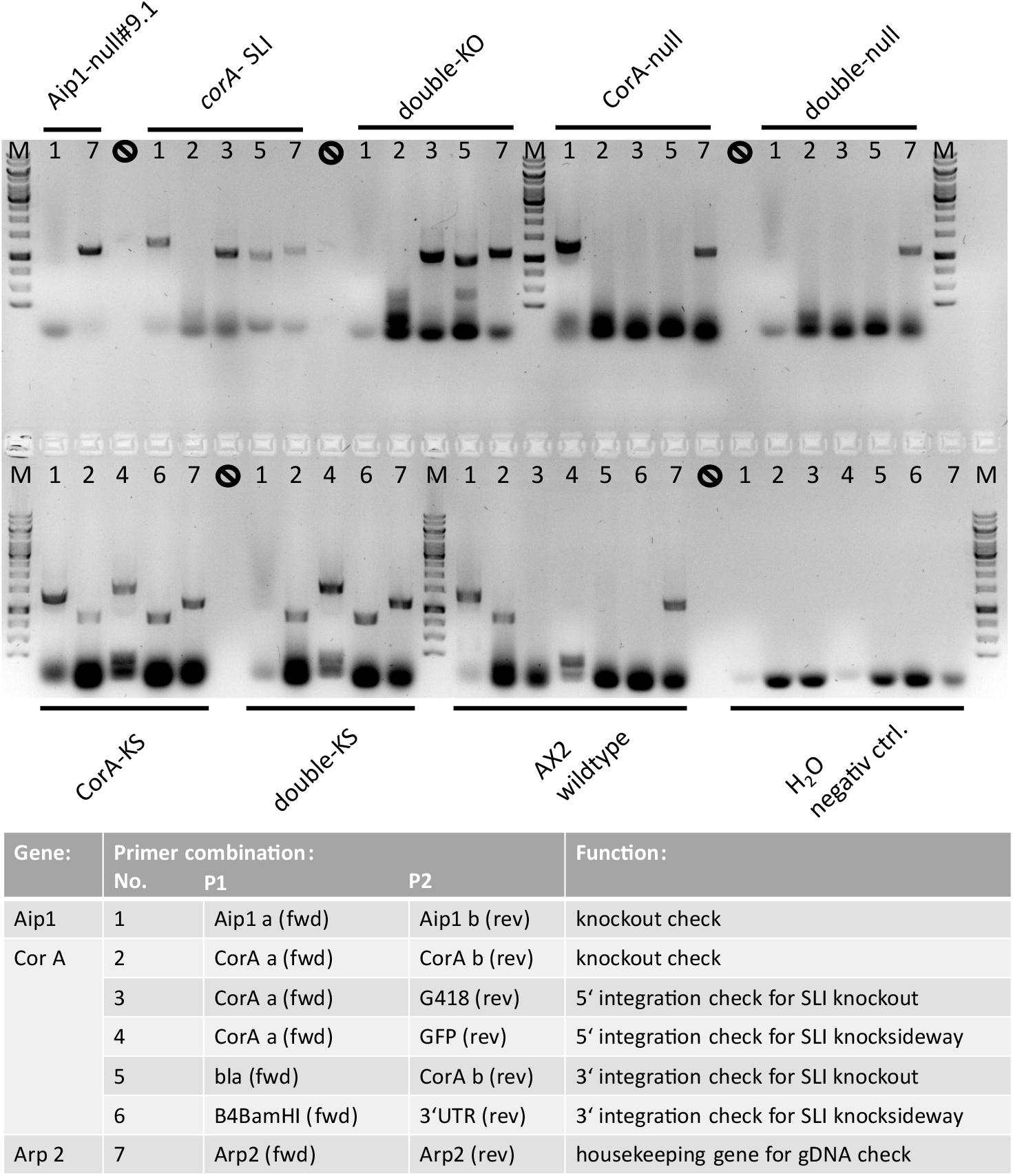
PCR for confirmation of aip1 and corA knockouts, as well as validation of knock-sideways constructs. On lysates of wild type AX2 cells and *corA* single and Aip1-null/*corA*-SLI double mutants knockouts and knock-sideways as well as loss-of-function cell lines; M: Marker, wild type AX2; Aip1-null#9.1 mutant [1]; *corA*-SLI: *corA* knockout done with SLI method; double-KO: *corA* knockout done with SLI method in Aip1-null mutant; CorA-null: classical gene disruption mutant of *corA* CorA- null HG1569#1 [3]; CorA-KS: Coronin-A knock-sideways done with SLI method; double-KS: Coronin-A knock-sideways done with SLI method in Aip1-null mutant.

**Figure S4.**
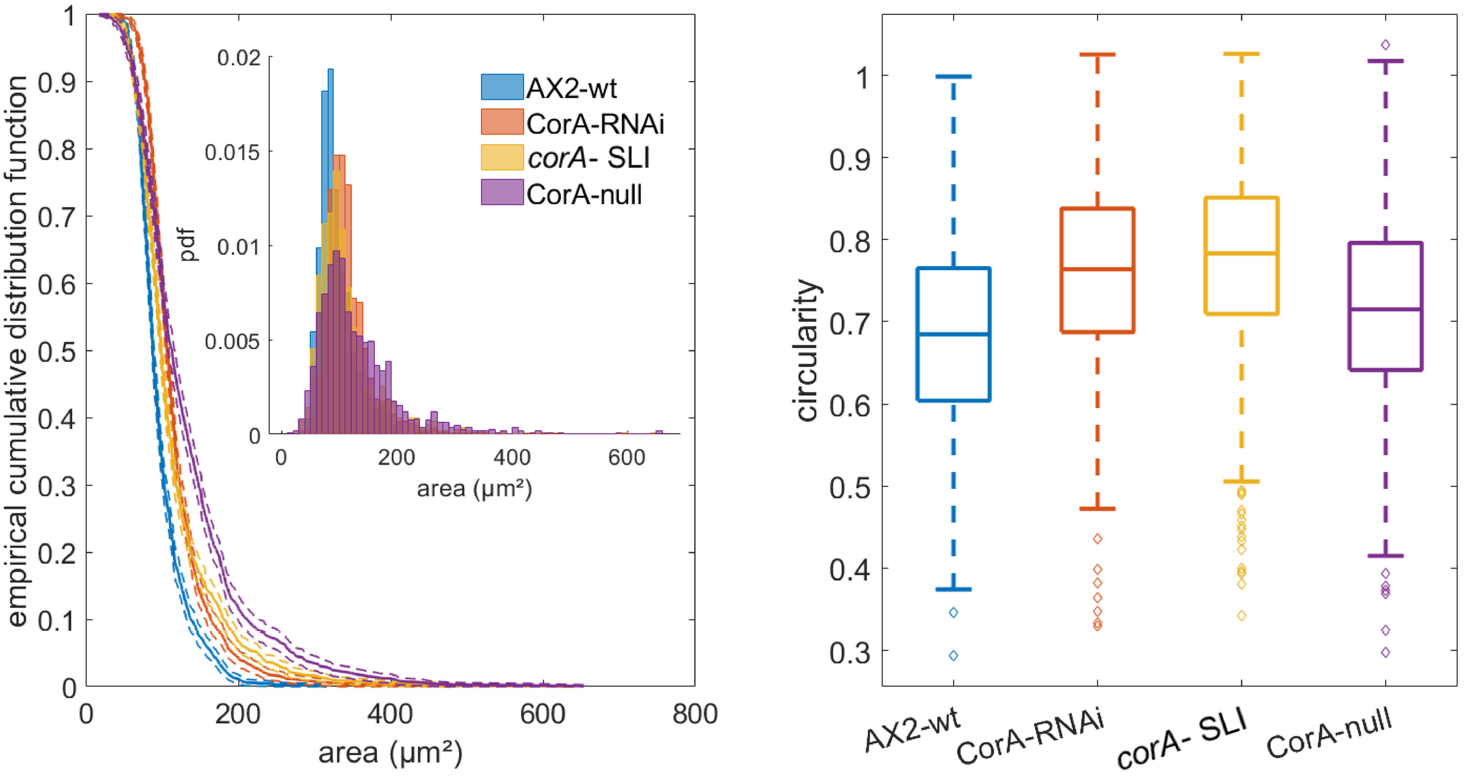
Illustration of cell size and cell circularity distribution in wild type AX2 cells, CorA knockdown by RNAi (CorA-RNAi), corA-SLI knockout done with SLI method and classical gene disruption mutant of corA CorA-null HG1569#1 ([3]); Left: empirical cumulative distribution function of the cell area; inset: histogram of the probability distribution of cell areas; right: circularity as the parameter for morphological differences shown as boxplots. For statistical analysis results of Kolmogorov-Smirnov test, see Tables S1-S2.

**Figure S5.**
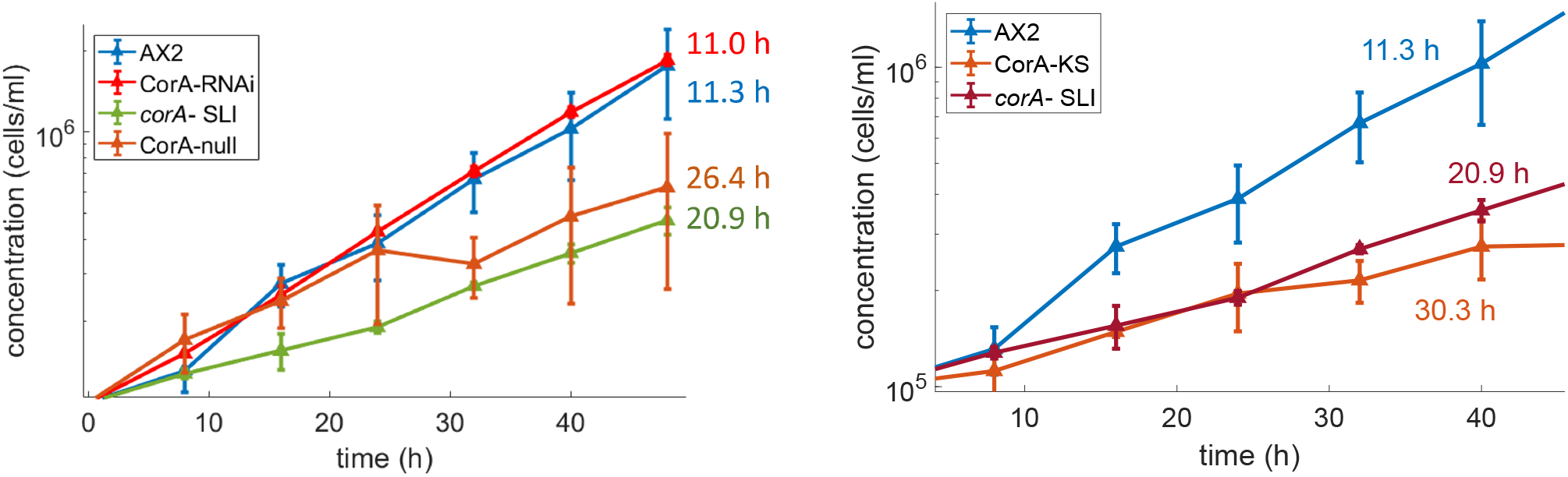
Growth curves in semi-log plot and doubling times. *Left:* growth of AX2 wild type, CorA-RNAi knockdown, *corA*-SLI knockout cells and CorA-null mutant cells (shaking culture in HL-5 medium) over 48 h. *Right:* data points show the mean cell concentration of AX2 wild type, CorA-KS, and *corA*-SLI knockout cells in shaking culture in HL-5 medium measured over a period of 48 h. In both cases, cell densities were determined as duplicates in 8 h intervals in two independent experiments, calculated by averaging the mean cell concentration determined for each experiment; error bars show the absolute difference between them (cf. Methods).

**Figure S6.**
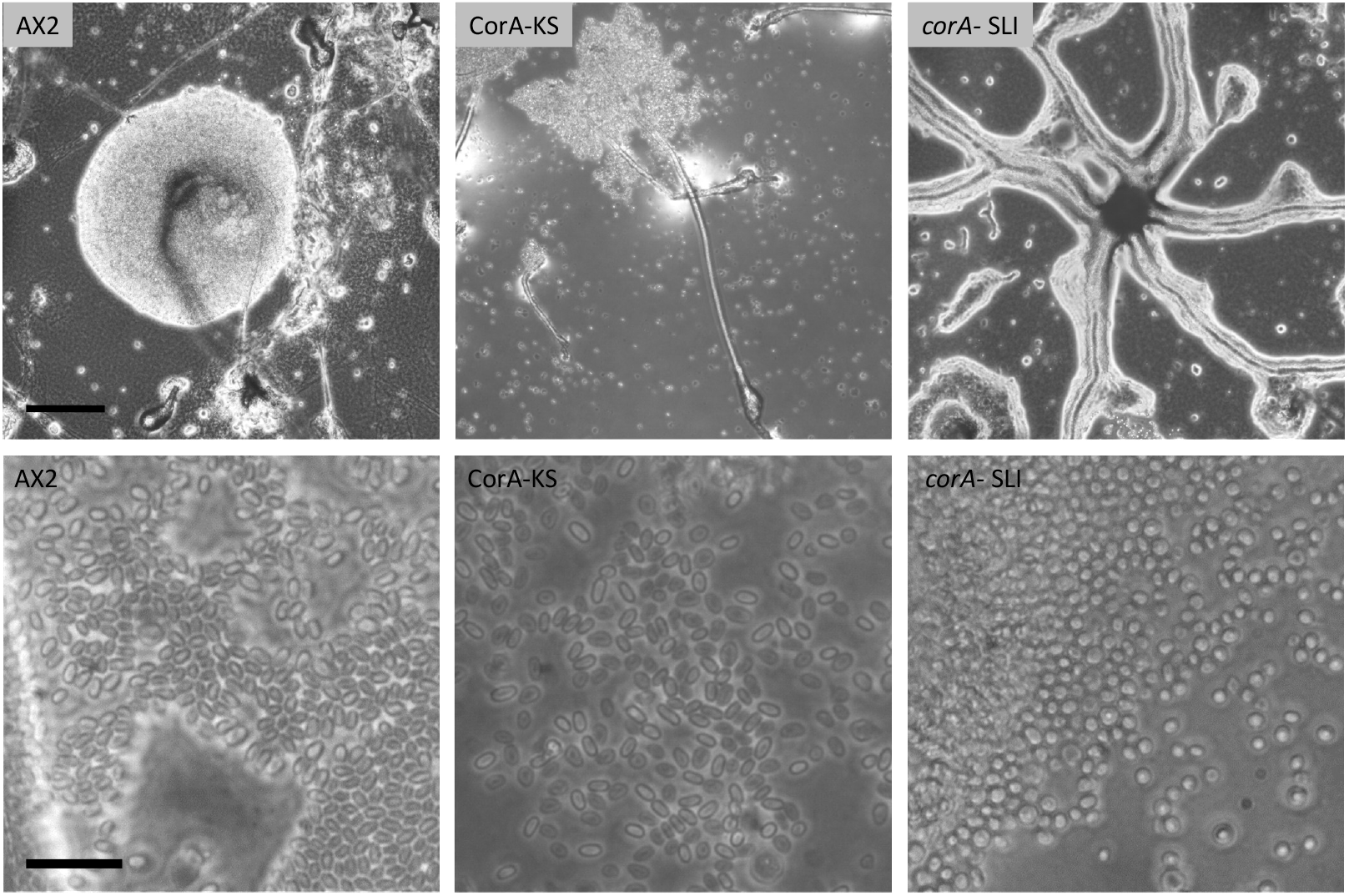
Influence of Coronin-A on development of fruiting bodies and spore formation. Representative examples are displayed of AX2 wild type, CorA-KS and *corA*-SLI knockout cell lines (from left to right), showing (upper) different stages of the aggregation process reached after 48 h; (lower) appearance of spores after 48 h; scale bars: 50 µm (upper panel) and 20 µm (lower panel).

**Figure S7.**
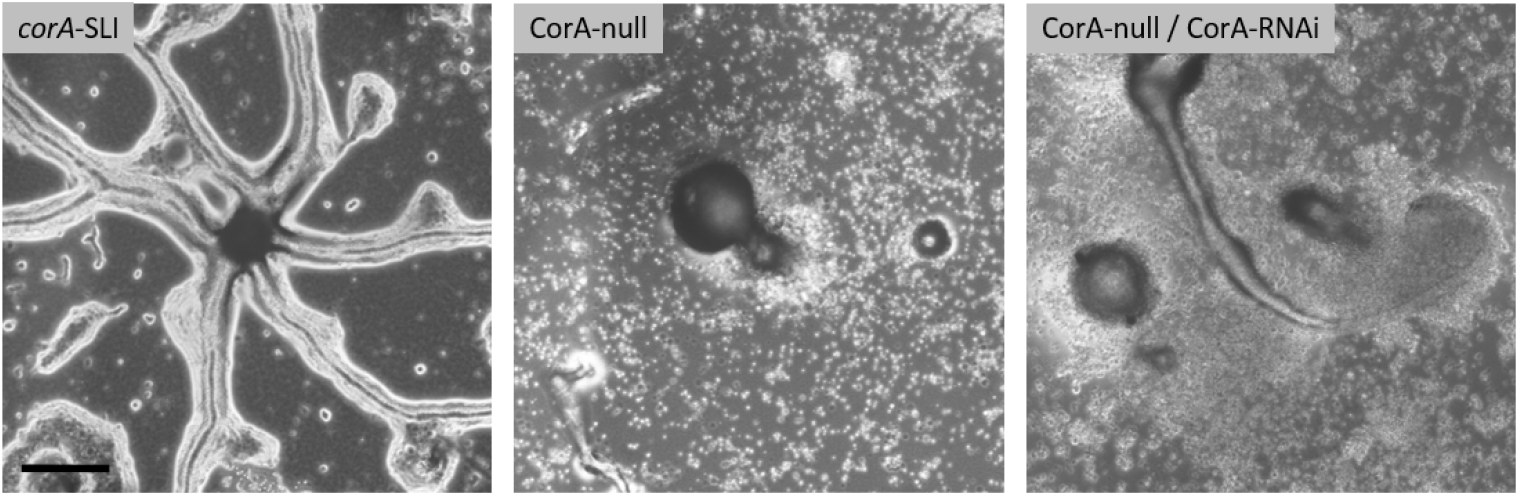
Influence of Coronin-A on development of fruiting bodies. Representative examples are displayed of *corA*-SLI knockout, CorA-null and CorA-null/CorA-RNAi cell lines (from left to right), showing different stages of the aggregation process reached after 48 h; scale bar: 50 µm.

**Figure S8.**
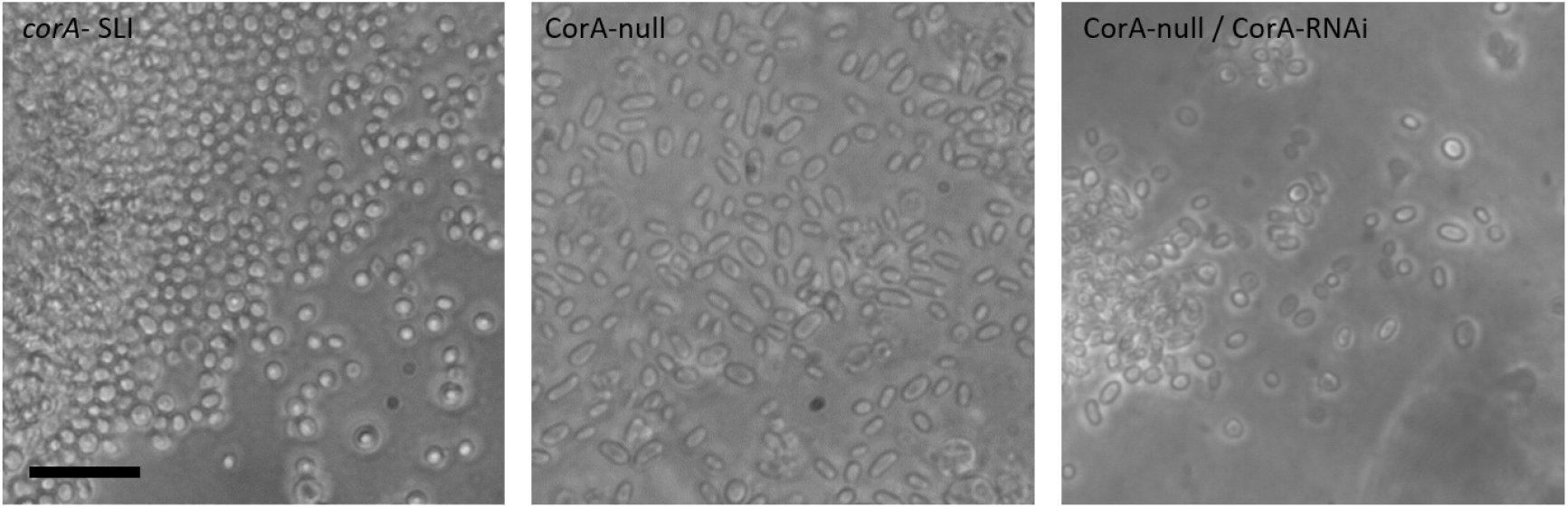
Influence of Coronin-A on spore formation. Representative examples are displayed of *corA*-SLI knockout, CorA-null and CorA-null/CorA-RNAi cell lines (from left to right), appearance of spores after 48 h; scale bar: 20 µm.

**Figure S9.**
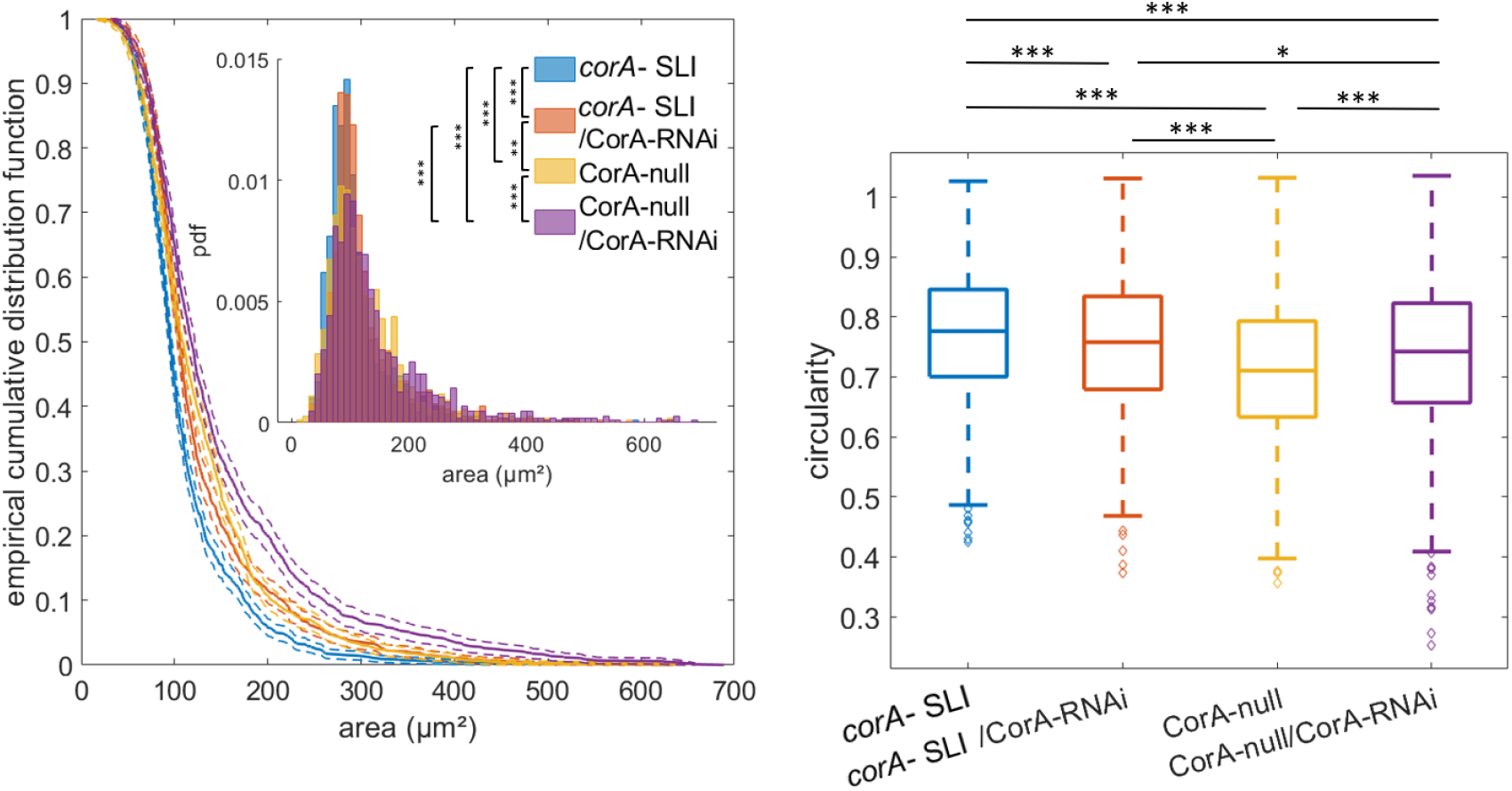
Illustration of cell size and cell circularity distribution in corA-SLI knockout done with SLI method and classical gene disruption mutant of corA CorA-null HG1569#1 [3] as well as both strains also expressing the CorA-RNAi; Left: empirical cumulative distribution function of the cell area; inset: histogram of the probability distribution of cell areas; right: circularity as the parameter for morphological differences shown as boxplots. For statistical analysis results of Kolmogorov-Smirnov test, see Tables S1-S2.

**Table S1.** Table of p-values calculated by Kolmogorov-Smirnov statistical test on the area of more than 1000 cells per strain as a parameter of morphology. For a more detailed strain description, see above and Table S3.

| strains | CorA-RNAi | CorA-OX | AX2-wt | AX2-wt (+) | aip1-/corA-RNAi | Aip1-null#9.1 | Aip1-null#9.1 (+) | CorA-null | CorA-KS | CorA-KS (+) | corA-SLI | CorA-null-RNAi | corA-SLI-RNAi | corA-SLI (+) | double-KO | double-KO (+) | double-KS | double-KS (+) |
| --- | --- | --- | --- | --- | --- | --- | --- | --- | --- | --- | --- | --- | --- | --- | --- | --- | --- | --- |
| CorA-OX | 0.000000 |  |  |  |  |  |  |  |  |  |  |  |  |  |  |  |  |  |
| AX2-wt | 0.000000 | 0.000009 |  |  |  |  |  |  |  |  |  |  |  |  |  |  |  |  |
| AX2-wt (+) | 0.005218 | 0.000000 | 0.000000 |  |  |  |  |  |  |  |  |  |  |  |  |  |  |  |
| aip1-/corA-RNAi | 0.015352 | 0.000000 | 0.000000 | 0.000000 |  |  |  |  |  |  |  |  |  |  |  |  |  |  |
| Aip1-null#9.1 | 0.000000 | 0.017924 | 0.000942 | 0.000000 | 0.000000 |  |  |  |  |  |  |  |  |  |  |  |  |  |
| Aip1-null#9.1 (+) | 0.000000 | 0.000000 | 0.000000 | 0.000000 | 0.000000 | 0.000000 |  |  |  |  |  |  |  |  |  |  |  |  |
| CorA-null | 0.000000 | 0.000000 | 0.000000 | 0.000000 | 0.000000 | 0.000000 | 0.001907 |  |  |  |  |  |  |  |  |  |  |  |
| CorA-KS | 0.071576 | 0.000000 | 0.000000 | 0.000034 | 0.160746 | 0.000000 | 0.000000 | 0.000000 |  |  |  |  |  |  |  |  |  |  |
| CorA-KS (+) | 0.100730 | 0.000000 | 0.000000 | 0.000011 | 0.238552 | 0.000000 | 0.000000 | 0.000000 | 0.230935 |  |  |  |  |  |  |  |  |  |
| corA-SLI | 0.000000 | 0.006364 | 0.000000 | 0.000000 | 0.000190 | 0.000000 | 0.000000 | 0.000000 | 0.000008 | 0.000000 |  |  |  |  |  |  |  |  |
| CorA-null-RNAi | 0.000000 | 0.000000 | 0.000000 | 0.000000 | 0.000000 | 0.000000 | 0.001606 | 0.000086 | 0.000000 | 0.000000 | 0.000000 |  |  |  |  |  |  |  |
| corA-SLI-RNAi | 0.000037 | 0.000000 | 0.000000 | 0.000000 | 0.000000 | 0.000000 | 0.000085 | 0.001155 | 0.000000 | 0.000003 | 0.000000 | 0.000000 |  |  |  |  |  |  |
| corA-SLI (+) | 0.000000 | 0.029995 | 0.000000 | 0.000000 | 0.000060 | 0.000003 | 0.000000 | 0.000000 | 0.000000 | 0.000000 | 0.221137 | 0.000000 | 0.000000 |  |  |  |  |  |
| double-KO | 0.000000 | 0.000000 | 0.000000 | 0.000000 | 0.000000 | 0.000000 | 0.000000 | 0.000000 | 0.000000 | 0.000000 | 0.009966 | 0.000000 | 0.000000 | 0.000029 |  |  |  |  |
| double-KO (+) | 0.000000 | 0.000000 | 0.000000 | 0.000000 | 0.000000 | 0.000000 | 0.000000 | 0.006255 | 0.000000 | 0.000000 | 0.000000 | 0.000000 | 0.000000 | 0.000000 | 0.000009 |  |  |  |
| double-KS | 0.000000 | 0.000000 | 0.000101 | 0.000000 | 0.000000 | 0.000000 | 0.000000 | 0.000000 | 0.000000 | 0.000000 | 0.000000 | 0.000000 | 0.000000 | 0.000000 | 0.000000 | 0.000000 |  |  |
| double-KS (+) | 0.000000 | 0.129648 | 0.026811 | 0.000000 | 0.000000 | 0.026228 | 0.000000 | 0.000000 | 0.000000 | 0.000000 | 0.000001 | 0.000000 | 0.000000 | 0.000116 | 0.000000 | 0.000000 | 0.000000 |  |
| double-null | 0.000613 | 0.000000 | 0.000000 | 0.000000 | 0.000000 | 0.000000 | 0.000075 | 0.000190 | 0.000007 | 0.000038 | 0.000025 | 0.000000 | 0.024002 | 0.000002 | 0.000078 | 0.001367 | 0.000000 | 0.000000 |

**Table S2.** Table of p-values calculated by Kolmogorov-Smirnov statistical test on the circularity of more than 1000 cells per strain as a parameter of morphology. For a more detailed strain description, see above and Table S3.

| strains | CorA-RNAi | CorA-OX | AX2-wt | AX2-wt (+) | aip1-/corA-RNAi | Aip1-null#9.1 | Aip1-null#9.1 (+) | CorA-null | CorA-KS | CorA-KS (+) | corA-SLI | CorA-null-RNAi | corA-SLI-RNAi | corA-SLI (+) | double-KO | double-KO (+) | double-KS | double-KS (+) |
| --- | --- | --- | --- | --- | --- | --- | --- | --- | --- | --- | --- | --- | --- | --- | --- | --- | --- | --- |
| CorA-OX | 0.000000 |  |  |  |  |  |  |  |  |  |  |  |  |  |  |  |  |  |
| AX2-wt | 0.000000 | 0.000000 |  |  |  |  |  |  |  |  |  |  |  |  |  |  |  |  |
| AX2-wt (+) | 0.000000 | 0.080346 | 0.000000 |  |  |  |  |  |  |  |  |  |  |  |  |  |  |  |
| aip1-/corA-RNAi | 0.005254 | 0.000000 | 0.000000 | 0.000000 |  |  |  |  |  |  |  |  |  |  |  |  |  |  |
| Aip1-null#9.1 | 0.000000 | 0.000000 | 0.000229 | 0.000000 | 0.000000 |  |  |  |  |  |  |  |  |  |  |  |  |  |
| Aip1-null#9.1 (+) | 0.000086 | 0.000000 | 0.000000 | 0.000000 | 0.163447 | 0.000000 |  |  |  |  |  |  |  |  |  |  |  |  |
| CorA-null | 0.000000 | 0.000000 | 0.000000 | 0.000000 | 0.000012 | 0.077402 | 0.000018 |  |  |  |  |  |  |  |  |  |  |  |
| CorA-KS | 0.045428 | 0.000000 | 0.000000 | 0.000000 | 0.000009 | 0.000000 | 0.000003 | 0.000000 |  |  |  |  |  |  |  |  |  |  |
| CorA-KS (+) | 0.000000 | 0.000000 | 0.000000 | 0.000000 | 0.000000 | 0.000000 | 0.000000 | 0.000000 | 0.000000 |  |  |  |  |  |  |  |  |  |
| corA-SLI | 0.001529 | 0.000000 | 0.000000 | 0.000000 | 0.000000 | 0.000000 | 0.000000 | 0.000000 | 0.060972 | 0.000000 |  |  |  |  |  |  |  |  |
| CorA-null-RNAi | 0.005478 | 0.000000 | 0.000000 | 0.000000 | 0.790136 | 0.000000 | 0.481291 | 0.000000 | 0.000071 | 0.000000 | 0.000000 |  |  |  |  |  |  |  |
| corA-SLI-RNAi | 0.977754 | 0.000000 | 0.000000 | 0.000000 | 0.008575 | 0.000000 | 0.000743 | 0.000000 | 0.013718 | 0.000000 | 0.000091 | 0.010979 |  |  |  |  |  |  |
| corA-SLI (+) | 0.000000 | 0.000177 | 0.000000 | 0.000004 | 0.000000 | 0.000000 | 0.000000 | 0.000000 | 0.000000 | 0.000000 | 0.000000 | 0.000000 | 0.000000 |  |  |  |  |  |
| double-KO | 0.001712 | 0.000000 | 0.000000 | 0.000000 | 0.000000 | 0.000000 | 0.000000 | 0.000000 | 0.227925 | 0.000000 | 0.010119 | 0.000002 | 0.000977 | 0.000000 |  |  |  |  |
| double-KO (+) | 0.000000 | 0.000000 | 0.000000 | 0.000000 | 0.000000 | 0.000000 | 0.000000 | 0.000000 | 0.000074 | 0.000000 | 0.000279 | 0.000000 | 0.000000 | 0.000463 | 0.003156 |  |  |  |
| double-KS | 0.000000 | 0.000000 | 0.000000 | 0.000000 | 0.000000 | 0.000000 | 0.000000 | 0.000000 | 0.000505 | 0.000000 | 0.000077 | 0.000000 | 0.000000 | 0.000127 | 0.010818 | 0.325295 |  |  |
| double-KS (+) | 0.000000 | 0.000000 | 0.000000 | 0.000000 | 0.000000 | 0.000000 | 0.000000 | 0.000000 | 0.309117 | 0.000000 | 0.000000 | 0.000000 | 0.000000 | 0.000000 | 0.000000 | 0.000000 | 0.000000 |  |
| double-null | 0.000000 | 0.000006 | 0.000000 | 0.001945 | 0.000000 | 0.000000 | 0.000000 | 0.000000 | 0.000000 | 0.000000 | 0.000000 | 0.000000 | 0.000000 | 0.008158 | 0.000000 | 0.000016 | 0.000039 | 0.000000 |

**Table S3.** The table summarizes the number of recorded cell tracks and the fraction of long tracks in motility assay experiments. The (+)-symbol indicates rapamycin treatment (final concentration: 10 µM). The total recording time was two hours; tracks shorter than ten minutes were considered to be too short for reliable statistical analysis and were discarded from the analysis; tracks with a length of 115-120 min are the most common.

| Cell line | Number of total tracks/cells | % of tracks between 230 and 240 frames |
| --- | --- | --- |
| **AX2** | 261 | 27.6 |
| **AX2 (+)** | 108 | 36.1 |
| **Aip1-null#9.1** (DAip1-, DBS0252637) | 251 | 35.1 |
| **Aip1-null#9.1** (DAip1-, DBS0252637) (+) | 143 | 24.5 |
| **CorA-null** HG1569#1 (DBS0236171) | 203 | 23.6 |
| **CorA/Aip1 double-null** (DBS0350851) | 192 | 38.5 |
| **corA-SLI** | 236 | 34.7 |
| **corA-SLI (+)** | 101 | 30.7 |
| **double-KO:** Aip1-null#9.1/*corA*-SLI | 227 | 44.5 |
| **double-KO:** Aip1-null#9.1/*corA*-SLI (+) | 119 | 38.7 |
| **CorA-RNAi** | 174 | 32.8 |
| **Aip1-null#9.1/CorA-RNAi** | 292 | 18.8 |
| **CorA-null** HG1569#1 /**CorA-RNAi** | 151 | 29.1 |
| **corA-SLI/CorA-RNAi** | 218 | 35.8 |
| **CorA-KS** | 88 | 42.0 |
| **CorA-KS (+)** | 104 | 33.7 |
| **Aip1-null/CorA-KS:** Aip1-null#9.1 + CorA-KS | 98 | 62,2 |
| **Aip1-null/CorA-KS (+):** Aip1-null#9.1 + CorA-KS | 157 | 39.5 |
| **CorA-OX** | 95 | 41.1 |
| **CorA-OX (+)** | 99 | 53.5 |

## Notes

### Competing Interest Statement

The authors have declared no competing interest.

